## Supplemental information for "An Sfi1-like centrin-interacting centriolar plaque protein affects nuclear microtubule homeostasis"

### Supplemental figures and tables

**Contents**


Figures S1 to S8

Tables S1 to S3

Movie captions S1 to S3


Supplemental movies can be accessed under:
https://www.dropbox.com/s/p14kyoarbj8ki7o/Supplemental%20movies.zip?dl=0

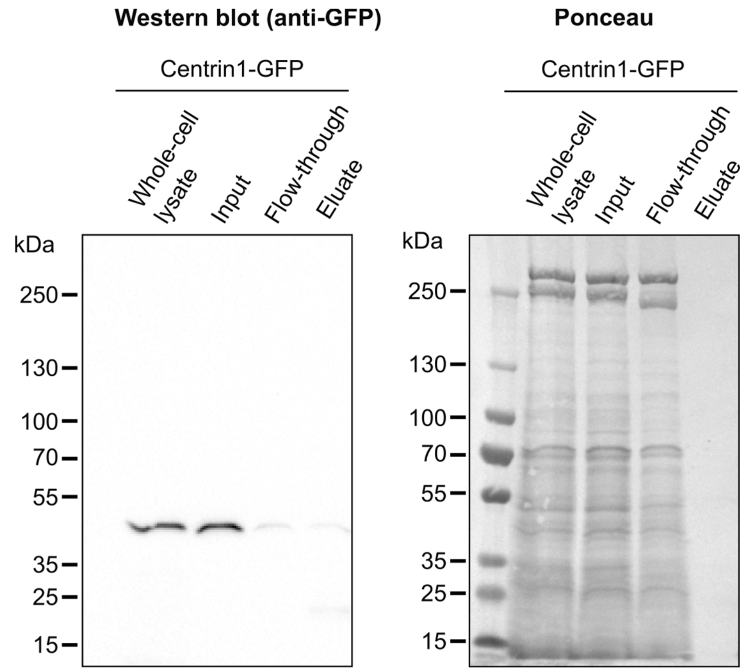


**Figure S1. Western blot analysis shows purification of PfCentrin1-GFP proteins via co-immunoprecipitation.** “Whole-cell lysate”, “input”, “flow-through” and “eluate” sample fractions were taken at different steps of PfCentrin1-GFP co-immunoprecipitation as described in the methods section. Size of the bands (circa 47 kDa) detected by the anti-GFP antibody in all lanes of the western blot corresponds to the molecular weight of PfCentrin1 (19.6 kDa) tagged with GFP (26.9 kDa). In the eluate fraction some degradation of PfCen1-GFP could be observed. Ponceau shows reduction of protein amount in eluate.

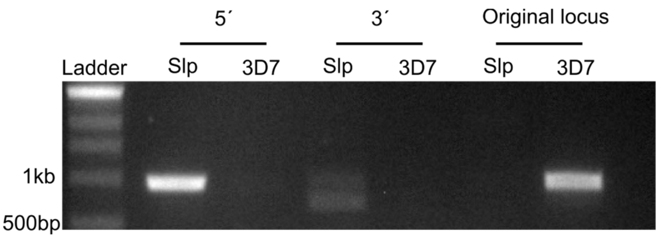


**Figure S2. PCR validation of genome integration of GFP-glmS-tag.** PCR of PfSlp-GFP transfected cells and 3D7 wildtype cells targeting 5´ and 3´ integrations to validate complete integration of the GFP-glmS tag to endogenous *PfSlp*. Control PCR targeting the unaltered locus of *PfSlp* in the 3D7 wild type strain confirms complete integration and modification of the endogenous locus.

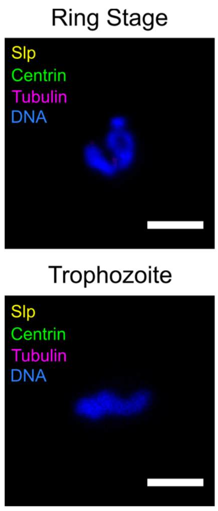


**Figure S3. PfSlp is not expressed before onset of schizogony.** Confocal microscopy images of immunofluorescence staining of ring stage parasites and trophozoites expressing endogenously tagged PfSlp-GFP using anti-centrin, anti-tubulin and anti-GFP antibodies. DNA stained with Hoechst. Maximum intensity projections are shown. Scale bars, 1.5 µm.

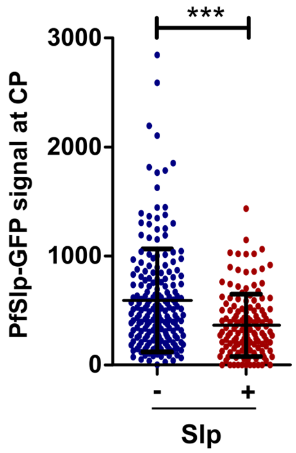


**Figure S4. GlcN-treatment causes reduction of PfSlp at the centriolar plaque.** Relative fluorescence signal intensity of PfSlp-GFP signal at the centriolar plaque was measured in +/-GlcN Slp schizont parasites with up to 10 nuclei, immunostained as in Fig. 1D.

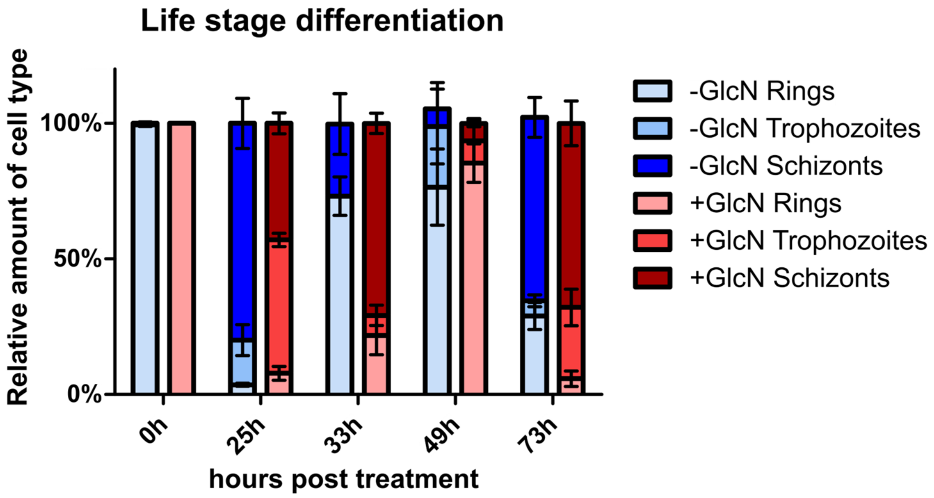


**Figure S5. PfSlp knock down delays progression from trophozoite to schizont stage.** Sorbitol synchronized PfSlp ring stage parasites +/-GlcN were fixed and giemsa stained at several timepoints throughout two life cycles. Asexual blood stages were determined by eye using light microscopy and the relative proportion of rings, trophozoites and schizonts was quantified. In GlcN treated cells, a delay of approximately 8 hours can be observed in the first generation after treatment, which is decreased in the second generation. Mean amounts with standard deviation were plotted using Prism GraphPad. N=3.

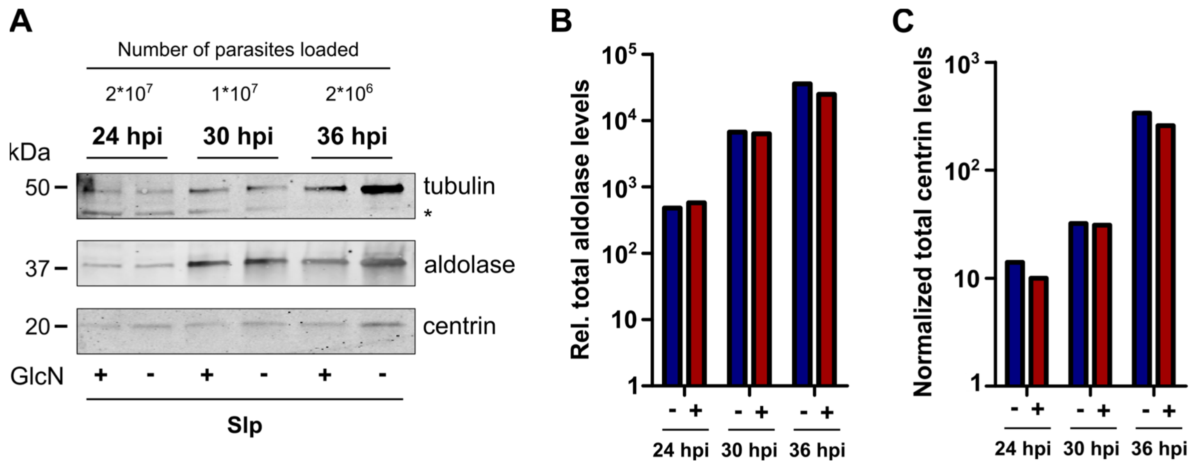


**Figure S6. Western blot analysis shows no increased tubulin abundance in Slp KD cells. A)** Synchronized Slp parasites +/- GlcN were harvested at 24, 30 or 36hpi, and SDS-PAGE of $2*10^7$, $1*10^7$ or $2*10^6$ parasites per lane respectively was performed. After blotting, the blot was cut at 25 kDa. The upper part was incubated with rabbit-anti-aldolase and mouse anti-tubulin primary antibodies, and anti-rabbit 680RD and anti-mouse 800CW secondaries, while the lower part was incubated with rabbit anti-Centrin3 primary and anti-rabbit 800CW secondary antibodies. Band at 50 kDa corresponds to the molecular weight of tubulin, a lower unspecific band is marked as (*). Band at 37 kDa and 20 kDa correspond to aldolase and centrin size respectively. Unspecific band was not included in measurement **B)** Quantification of aldolase protein signal in A) after correcting for equal parasite number. **C)** Quantification of centrin protein level in A) after correcting to equal parasite number, normalized to aldolase signal.

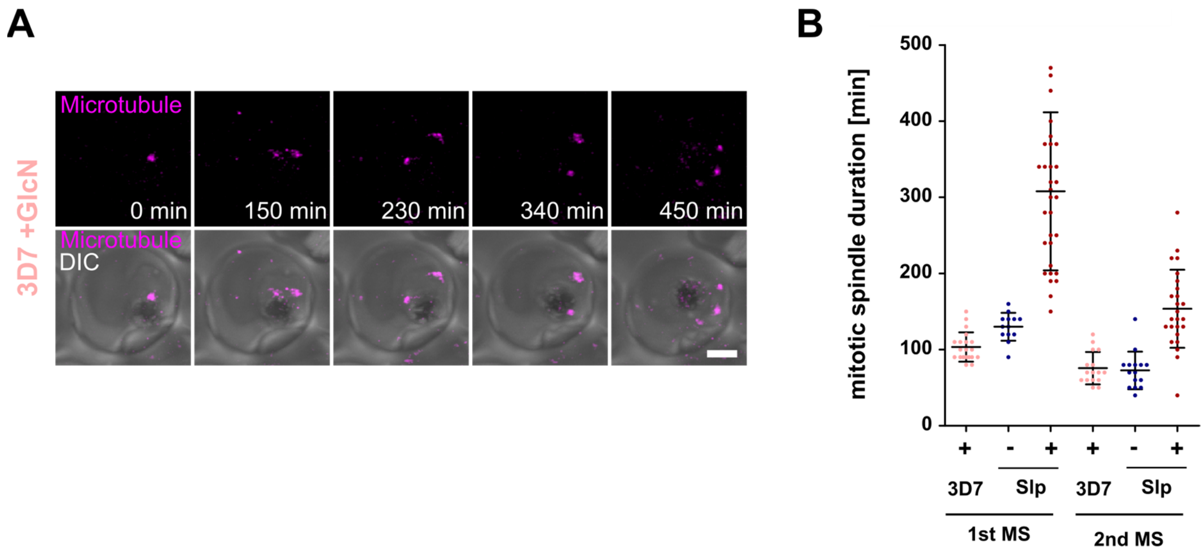


**Figure S7. GlcN treatment or *glmS* tagging alone does not strongly delay mitotic spindle extension.** A) Time-lapse confocal imaging of 3D7 parasites +GlcN undergoing schizogony stained with SPY555-Tubulin (magenta). Maximum intensity projections are shown. Scale bar, 1.5 µm. B) Quantification of first (3D7 (n=20) and second (3D7 (n=16)) mitotic spindle stage duration using live-cell movies of tubulin-stained parasites that complete spindle extension.

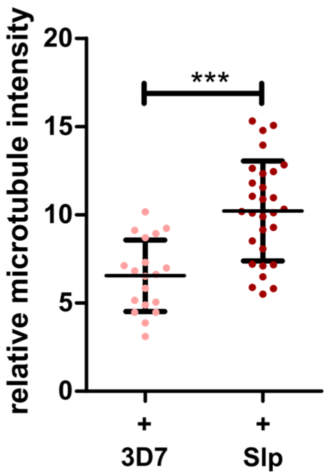


**Figure S8. Microtubule levels are already increased in the first mitotic spindle of PfSlp knock down parasites.** Movies of SPY555-Tubulin labelled 3D7 +GlcN and Slp +GlcN treated cells acquired in the same imaging session using identic settings. For quantification the first timeframe containing showing a defined mitotic spindle were selected and relative microtubule signal intensity was quantified using average intensity projections of image slices containing the spindle microtubule signal.

| Gene ID | Identified Proteins | M.W. | Peptide count | GFP control pull down (Balestra et al. 2021) |
|---|---|---|---|---|
| **PF3D7_0107000** | **Centrin-1** | **20 kDa** | **24** | **not identified in GFP control** |
| PF3D7_1357000 | Elongation factor 1-alpha | 49 kDa | 5 | identified in GFP control |
| PF3D7_1462800 | Glyceraldehyde-3-phosphate dehydrogenase | 37 kDa | 5 | identified in GFP control |
| PF3D7_0818900 | Cluster of Heat shock protein 70 | 74 kDa | 5 | identified in GFP control |
| **PF3D7_1027700** | **Centrin-3** | **21 kDa** | **4** | **not identified in GFP control** |
| PF3D7_1117700 | GTP-binding nuclear protein | 25 kDa | 3 | identified in GFP control |
| **PF3D7_0710000** | **Conserved protein, unknown function** | **407 kDa** | **3** | **not identified in GFP control** |
| PF3D7_0610400 | Histone H3 | 15 kDa | 2 | identified in GFP control |
| PF3D7_0719600 | 60S ribosomal protein L11a, putative | 20 kDa | 2 | identified in GFP control |
| PF3D7_0708400 | Heat shock protein 90 | 86 kDa | 2 | identified in GFP control |
| PF3D7_1116800 | Heat shock protein 101 | 103 kDa | 2 | identified in GFP control |

**Supplemental Table S1. Mass spectrometry analysis of Centrin Co-IP reveals two specific interaction partners.** All proteins identified from the non-crosslinked PfCentrin1-GFP co-immunoprecipitation samples are listed with peptide counts. Only Centrin-1, Centrin-3, and PfSlp (highlighted in green) were not detectable in a GFP only control pull down done with the same protocol used in a previous study by Balestra et al. 2021.

| Primer designation | Sequence |
|---|---|
| 1: Slp cDNA fw 1 | GAAGACGATGTTGAGGAGGGG |
| 2: Slp cDNA rev 1 | TCGTCGTTATGGACATCCTCT |
| 3:Slp cDNA fw 2 | AGGTGAAAGTATAAGCGGTCAGG |
| 4: Slp cDNA rev 2 | CCCCTCCTCAACATCGTCTT |
| 5: Serine tRNA ligase fw | AAGTAGCAGGTCATCGTGGTT |
| 6: Serine tRNA ligase rev | TTCGGCACATTCTTCCATAA |
| 7: gDNA 5´ integration fw | CACCACATCTTCATAACTCTTCAGG |
| 8: gDNA 5´ integration rev | GCATCACCTTCACCCTCTCC |
| 9: gDNA 3´ int fw | GAGCGGATAACAATTTCAC |
| 10: gDNA 3´ int rev | CAAAACATGTTTACATTATTGACAAGG |
| 7: gDNA original locus fw | CACCACATCTTCATAACTCTTCAGG |
| 10: gDNA original locus rev | CAAAACATGTTTACATTATTGACAAGG |

**Supplemental Table S2. List of primers used in this study.**

| Antibody | Species | Dilution | Source |
|---|---|---|---|
| anti-alpha-tubulin B-5-1-2, monoclonal | mouse | 1:500 | Sigma |
| Anti-tubulin [YOL1/34] | rat | 1:200 | Abcam |
| anti-PfCentrin3, polyclonal | rabbit | 1:500 (1:1000 for WB) | Simon et al., 2021 |
| Anti-GFP 3E6, monoclonal | mouse | 1:200 | Thermo |
| Anti-EGFP [F56-6A1.2.3] | mouse | 1:50 | Abcam |
| Anti GFP (11814460001, Western blot) | mouse | 1:2000 | Roche |
| HRP Anti- PfAldolase antibody (ab38905) | rabbit | 1:1000 | Abcam |
| anti-rat-Alexa 488 | Goat | 1:1000* | Sigma |
| anti-mouse-Alexa 568 | Goat | 1:1000 | Sigma |
| anti-mouse-Atto647 | Goat | 1:1000* | Sigma |
| anti-rabbit-Atto594 | Goat | 1:1000* | Sigma |
| anti-rabbit-Atto647 | Goat | 1:1000 | Sigma |
| Anti-mouse IgGHRP (A5278) | Goat | 1:3000 | Sigma |
| IRDye® 800CW anti-Mouse IgG (H + L) | Goat | 1:10.000 | LI-COR |
| IRDye® 680RD anti-Rabbit IgG (H + L) | Goat | 1:10.000 | LI-COR |
| IRDye® 800CW anti-Rabbit IgG (H + L) | Goat | 1:10.000 | LI-COR |
| **Dye** | | **Dilution** | **Source** |
| SPY555-Tubulin (SC203) | - | 1:2000 | Spirochrome |
| Hoechst33342 | - | 1:1000 | Thermo |

**Supplemental Table S3: Antibodies and dyes used in this study.** Starred (*) indicates dilutions for IFAs imaged by confocal microscopy; for STED, those antibodies were used at 1:200.

**Movie S1. Normal mitotic spindle formation and extension dynamics in Slp parasite.** Super-resolution time-lapse confocal microscopy of Slp blood stage parasite -GlcN labelled with microtubule live cell dye SPY555-Tubulin (magenta) and acquisition of transmission mode (grey) undergoing the first few mitotic divisions. Images were acquired with Zeiss Airyscan detector and processed for improved resolution. Maximum projections are shown. Image dimensions are 10 x 10 µm.

**Movie S2. Normal mitotic spindle formation and extension dynamics in wild type parasite.** Super-resolution time-lapse confocal microscopy of 3D7 wild type blood stage parasite +GlcN labelled with microtubule live cell dye SPY555-Tubulin (magenta) and acquisition of transmission mode (grey) undergoing the first few mitotic divisions. Images were acquired with Zeiss Airyscan detector and processed for improved resolution. Maximum projections are shown. Image dimensions are 10 x 10 µm.

**Movie S3. Mitotic spindle extension is strongly delayed in PfSlp knock down.** Super-resolution time-lapse confocal microscopy of Slp blood stage parasite +GlcN labelled with microtubule live cell dye SPY555-Tubulin (magenta) and acquisition of transmission mode (grey) attempting the first mitotic spindle extension. Images were acquired with Zeiss Airyscan detector and processed for improved resolution. Maximum projections are shown. Image dimensions are 10 x 10 µm.
